# cAMP in budding yeast: a messenger for sucrose metabolism?

**DOI:** 10.1101/2023.12.15.571809

**Authors:** Dennis Botman, Sineka Kanagasabapathi, Mila I. Rep, Kelly van Rossum, Evelina Tutucci, Bas Teusink

**Affiliations:** Systems Biology Lab, AIMMS/ALIFE, Vrije Universiteit Amsterdam, 1081 HV, Amsterdam, The Netherlands

**Author notes:** Correspondence should be addressed to Dennis Botman or Bas Teusink. These authors contributed equally to this work (Dennis Botman, Sineka Kanagasabapathi).

**Keywords:** Saccharomyces cerevisiae, metabolism, cAMP, sucrose, glucose, signalling, G-protein coupled receptor

## Abstract

*S. cerevisiae* (or budding yeast) is an important micro-organism for sucrose-based fermentation in biotechnology. Yet, it is largely unknown how budding yeast adapts to sucrose transitions. Sucrose can only be metabolized when the invertase or the maltose machinery are expressed and we propose that the Gpr1p receptor signals extracellular sucrose availability via the cAMP peak to adapt cells accordingly. A transition to sucrose or glucose gave a transient cAMP peak which was maximally induced for sucrose. When transitioned to sucrose, cAMP signalling mutants showed an impaired cAMP peak together with a lower growth rate, a longer lag phase and a higher yield compared to a glucose transition. These effects were not caused by altered activity or expression of enzymes involved in sucrose metabolism and imply a more general metabolic adaptation defect. Basal cAMP levels were comparable among the mutant strains, suggesting that the transient cAMP peak is required to adapt cells correctly to sucrose. We propose that the short-term dynamics of the cAMP signalling cascade detects long-term extracellular sucrose availability and speculate that its function is to maintain a fermentative phenotype at continuously low glucose and fructose concentrations.

## Introduction

To stay competitive most life on earth needs to adapt to continuously changing environments. Most eukaryotes, including budding yeast (*Saccharomyces cerevisiae)* studied here, rely on sensing changes in available nutrients and adapting gene expression accordingly. Budding yeast resides in nature on fruit and on tree barks where it encounters sucrose, the end-product of plant photosynthesis^1^. In industry, sucrose is a widely used fermentable sugar to produce both food and beverages, but also pharmaceuticals and other chemicals^2–5^.

Although sucrose-based fermentation using budding yeast is widely used, remarkably little is known about its physiology under this condition. The limited data available show that laboratory yeast strains (i.e. Cen.PK) grow slower on sucrose (0.38 h^-1^) compared to glucose (0.41 h^-1^), while other wild strains show a growth rate up to 28% higher on sucrose than on glucose^6,7^. The mechanisms underlying these differences are unknown and therefore, a better understanding of the sensing and adaptation of budding yeast to sucrose would benefit biotechnological processes.

Sucrose is a disaccharide that needs to be hydrolyzed to glucose and fructose. This happens both extracellularly by *SUC2*, and intracellularly^8^. For intracellular hydrolysis, sucrose is transported by maltose transporters *MAL11, MAL12, MAL22, MAL31, MAL32, MPH2 and MPH3* where transport by Mal11p is probably the main mode^9^, whereafter it is hydrolyzed by the isomaltoses *IMA1-5*^9–11^. These enzymes have a K_m_ ranging from 116 to 191 mM and a k_cat_ 25-55 s^-1^ with the exception of IMA5 with a k_cat_ of 4 s^-1 12^. However, how much a yeast cell uses intracellular sucrose hydrolyses varies between strains^10^. In some strains, sucrose metabolism occurs mostly extracellularly^13–17^ and diffusion and consumption by (competing^18^) cells potentially makes the resultant concentrations of extracellular glucose and fructose relatively low. To stay fit, cells should signal the sucrose availability in the extracellular environment and steer themselves to a maximal growth state. For strains that metabolize sucrose intracellularly, extracellular detection of sucrose is also necessary to express the appropriate transporters and enzymes^11,19^.

We hypothesized that the cAMP-PKA signalling pathway scouts specifically for sucrose availability in the environment (Fig. 1A). Activation of cAMP-PKA occurs when respiring or non-growing cells are transitioned to an environment containing a fermentable carbon source such as glucose, fructose or the disaccharide sucrose. This transition gives a distinctive cAMP peak followed by an elevated baseline (Fig. 1B). cAMP activates PKA, a protein kinase that changes the metabolic program from a slow or no growth to a fermentative growth state^20^. cAMP synthesis is affected via metabolism of sugars and sensing of sugars by the G-protein coupled receptor Gpr1p^21–24^. The exact purpose of the Gpr1p receptor is enigmatic, as *Gpr1* deletion mutants show no clear defects in fitness when grown on glucose^24^. Since the cAMP-PKA signalling cascade reacts to sucrose and it is mostly metabolized outside the cell, we hypothesise that the Gpr1p receptor cascade has evolved to detect sucrose in the extracellular environment and adapt the cells properly to its availability.

**Figure 1.**
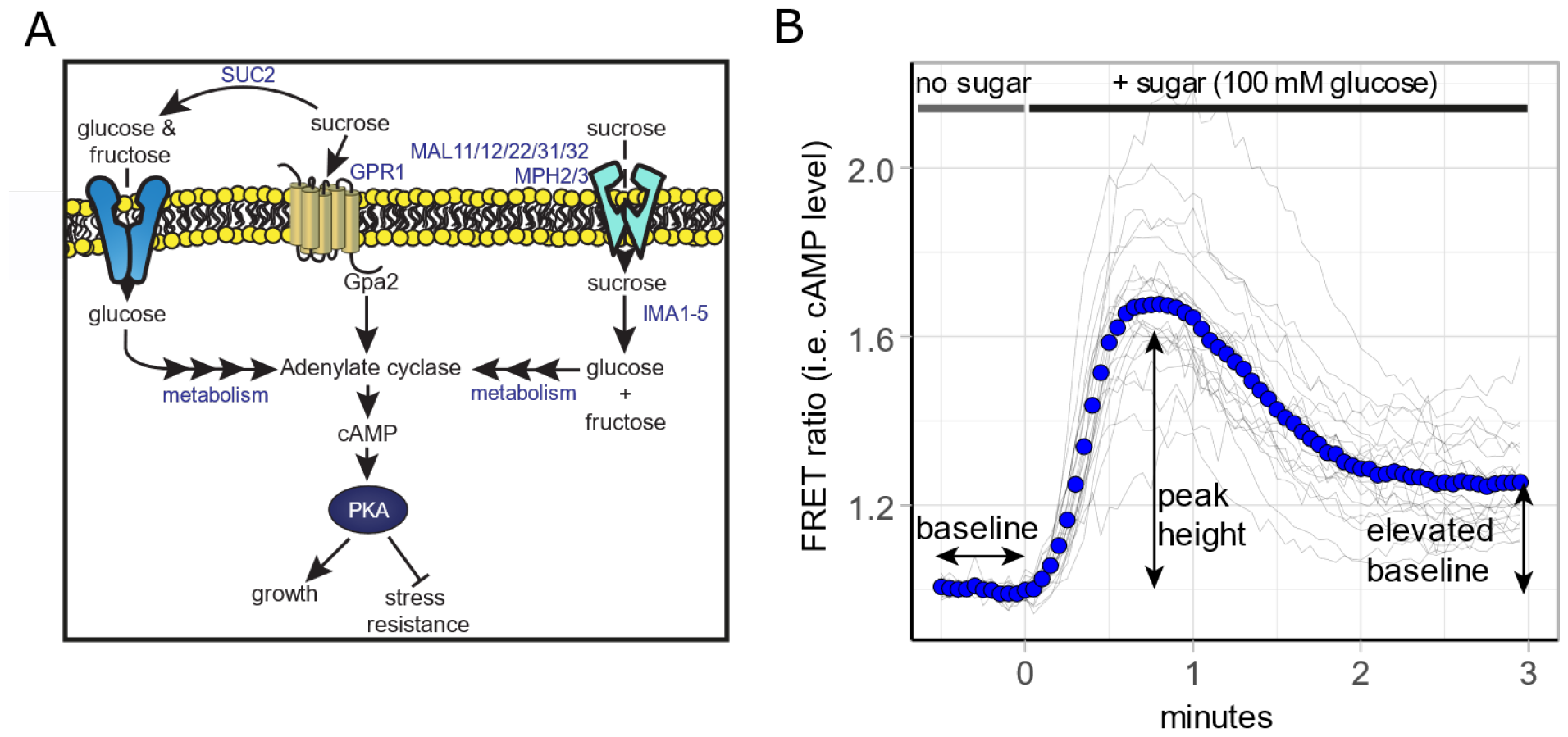
cAMP-PKA signalling in budding yeast. A) Simplified network for glucose and sucrose metabolism and signalling. B) A typical short-term cAMP response of yeast grown on a respiring substrate to a 100 mM glucose addition. Blue dots show mean response, grey lines show single-cell traces.

Currently, little evidence exists to support this hypothesis. Some studies indeed imply that cAMP regulates *SUC2* expression^25–29^, but these effects are strain dependent. For *SUC2*, some studies show a reduced invertase activity in cAMP-PKA signalling mutants^30,31^. Like *SUC2*, the *MALx2* and *IMA* genes are subject to glucose repression, where the cAMP peak is involved^9,11,32^. Furthermore, the *IMA2, IMA3* and *IMA4* genes contain a STRE motif which is regulated by Msn2/Msn4, two main transcription factors inhibited by cAMP-PKA signalling cascades^11,33^. Finally, some studies show that the industrial yeast strain Cen.PK (which has the *Cyr1*^*K1876M*^ mutation, affecting the cAMP peak^34,35^) has impaired growth on sucrose compared to other yeast strains but little difference is found between various components of cAMP signalling cascade during steady-state growth^7,34^. We hypothesise that the short-termed transient cAMP peak is essential to reach the correct growth state on sucrose and that this is not observable during steady-state measurements. Here, we explore and provide evidence that supports our hypothesis.

## Material and methods

### Yeast growth

Strains used in this study are listed in table 1. Cells were grown in YNB medium (Sigma Aldrich, St. Louis, MO, USA), containing 1% ethanol (VWR International, Radnor, PA, USA), 20 mg/L adenine hemisulfate (Sigma-Aldrich), 20 mg/L L-tryptophan (Sigma-Aldrich), 20 mg/L L-histidine (Sigma Aldrich), 60 mg/L L-leucine (SERVA Electrophoresis GmbH, Heidelberg, Germany), a carbon source (100 mM or 1% v/v), and 20 mg/L uracil (Sigma Aldrich) for strains not carrying a plasmid. The next day, cells were diluted and grown to an OD_600_ of 0.1-2 with at least 5 divisions. Next, cells were either used for microscopy or growth assays.

**Table 1.**
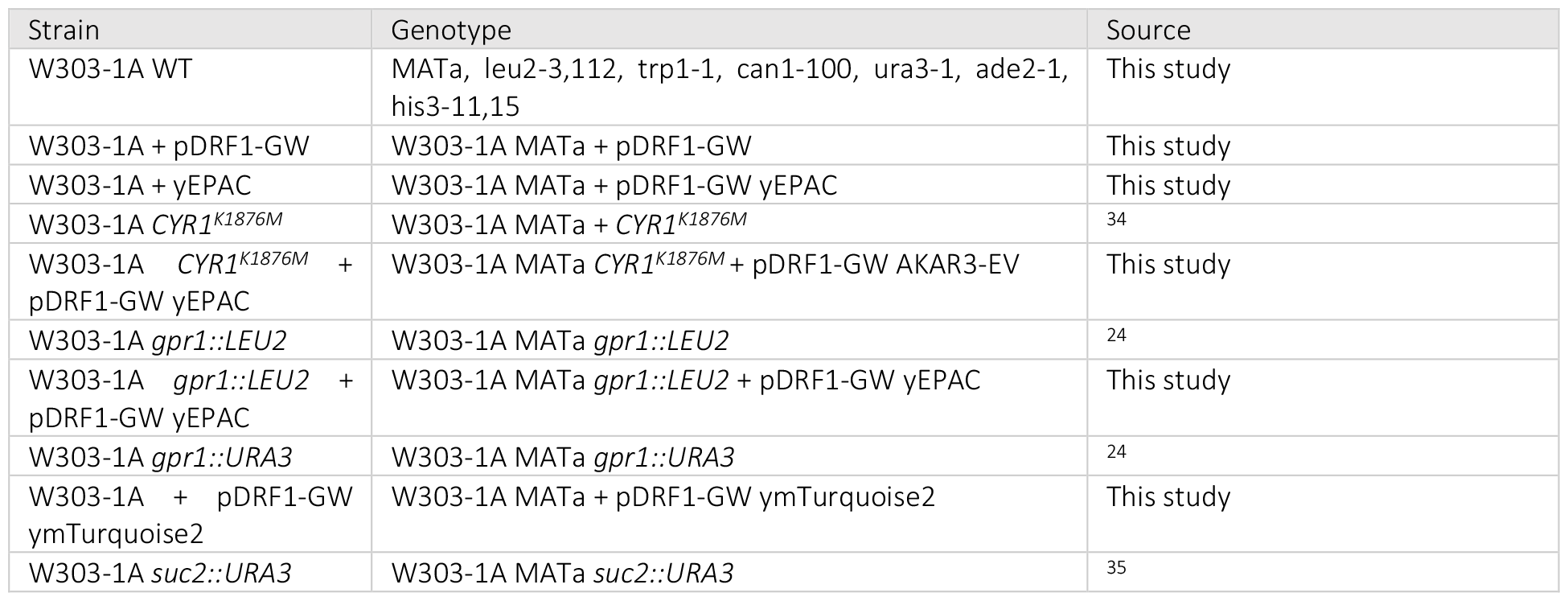
Used strains in this study.

| Strain | Genotype | Source |
| --- | --- | --- |
| W303-1A WT | MATa, leu2-3,112, trp1-1, can1-100, ura3-1, ade2-1, his3-11,15 | This study |
| W303-1A + pDRF1-GW | W303-1A MATa + pDRF1-GW | This study |
| W303-1A + yEPAC | W303-1A MATa + pDRF1-GW yEPAC | This study |
| W303-1A *CYR1*<sup>K1876M</sup> | W303-1A MATa + *CYR1*<sup>K1876M</sup> | 34 |
| W303-1A *CYR1*<sup>K1876M</sup> + pDRF1-GW yEPAC | W303-1A MATa *CYR1*<sup>K1876M</sup> + pDRF1-GW AKAR3-EV | This study |
| W303-1A *gpr1::LEU2* | W303-1A MATa *gpr1::LEU2* | 24 |
| W303-1A *gpr1::LEU2* + pDRF1-GW yEPAC | W303-1A MATa *gpr1::LEU2* + pDRF1-GW yEPAC | This study |
| W303-1A *gpr1::URA3* | W303-1A MATa *gpr1::URA3* | 24 |
| W303-1A + pDRF1-GW ymTurquoise2 | W303-1A MATa + pDRF1-GW ymTurquoise2 | This study |
| W303-1A *suc2::URA3* | W303-1A MATa *suc2::URA3* | 35 |

### ConA coverslips

Concanavalin A(ConA) coverslips were made as described by Hansen et al., 2015^54^. Next, ConA was diluted to a final concentration of 200 μg/mL and coverslips were covered with ConA. Finally, the coverslips were dried overnight in 6 wells plates.

### Microscopy

The grown cells were transferred to 6 wells plate containing ConA-coated coverslips. To visualize a coverslip, the coverslip was mounted in an Attofluor cell chamber (ThermoFisher Scientific, Waltham, MA) and 1 mL of medium was added. Samples were visualized with a Nikon Ti-eclipse microscope (Nikon, Minato, Tokyo, Japan) at 30°C equipped with a TuCam system (Andor, Belfast, Northern Ireland) and 2 Andor Zyla 5.5 sCMOS Cameras (Andor) and a SOLA 6-LCR-SB light source (Lumencor, Beaverton, OR). yEPAC was visualized using 438/24 nm excitation and emission was passed through a 458 nm long-pass (LP) dichroic mirror. Acceptor and donor emission was obtained using the TuCam system containing a 552 nm LP dichroic, 483/32 nm and 593/40 nm filter set. All filters are from Semrock, Lake Forest, IL. In the time-lapse experiments, a baseline was recorded after which the compound was added from a 10x concentration to a final concentration of 1x.

For analysis, cells were segmented using a Fiji^36^ and an in-house script which reduces drift using the image stabilizer plugin^55^, corrects for background correction and performs segmentation using the Weka Segmentation plugin^56,57^. Mean fluorescence was calculated for each cell per frame and results were analysed using R 4.2.2. For all cells, 11% bleed-trough correction was performed and the FRET ratio (i.e. the CFP : bleed-trough-corrected RFP ratio) was calculated. Finally, baseline normalization was performed for time-lapse data. For the dose-response fit, maximal FRET levels after the sugar additions were fitted against the final sugar concentration according equation 1, using the nls function in R, with [sugar] as the sugar concentration in mM, max as the maximal normalized FRET value and K_0.5_ as the sugar concentration giving 50% of the maximal response.

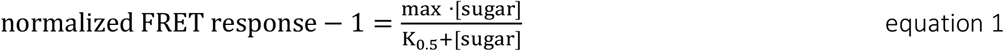

### Growth assays

Cells grown as aforementioned were washed twice (by centrifuging at 3300 g for 3 minutes and resuspending in YNB medium without carbon source) and resuspended in YNB medium without carbon source to an OD_600_ of 1 and 20 μL was transferred to a 48-well plate containing 480 μL YNB medium containing 10 mM or 1% of a carbon source. For growth assays without washing, grown cells were diluted or concentrated to an OD_600_ of 1 and were transferred to a 48-wells plate as described. Cells were grown in either a CLARIOstar (all growth assays except for Fig. S2) or Fluostar plate reader (for Fig. S2, BMG LABTECH, Ortenberg, Germany) at 30°C and 700 rpm orbital shaking and OD_600_ was measured every 5 minutes.

For analysis, the OD_600_ values were ln-transformed a moving average was performed after which a linear regression was fitted in a sliding window. The slope of this linear regression fit is the growth rate in the window. Maximal growth rates from these linear regression fits were obtained for each well. The time point at which the maximal growth rate was obtained was used as approximation for the time of the lag phase. Yield was calculated as the median OD obtained between 35 and 40 hours of growth.

### Invertase assay

Cells were grown as described with 1% ethanol as carbon source. The first sample was obtained from this culture. Next, cells were washed in YNB without carbon source twice (as described in the growth assays section) and cells were resuspended in YNB medium containing the appropriate supplements (as described in the yeast growth section) and 100 mM sucrose to an OD_600_ of 0.04. Afterwards, samples were taken at various time points and invertase activity was measured using the Invertase Activity Assay Kit (Merck, Darmstadt, Germany) according to the manufacturer’s protocol. In brief, 40 μL of cells (undiluted for the first 2 time points, 10x diluted for the third time point and 31x diluted for the last 2 time points) were incubated with 5 μL of 2 M sucrose in YNB for 20 minutes at 30 ° C, 90 μL reaction buffer was added and incubated for 30 minutes after which OD_570_ was measured. Finally, OD_570_ values were normalized for OD_600_ values (cell density of the culture) and normalized to the second time point of the experiment.

### Flow cytometry

Cells were grown as described with YNB medium containing either 100 mM glucose or 100 mM sucrose. Next, samples were measured using a CytoFLEX S Flow Cytometer (Beckman Coulter, Brea, CA, United States). Cells were excited using a 405 nm laser and fluorescence was recorded using avalanche photodiodes at 470/20 and 610/20 nm. Events with a saturating forward or side scatter were filtered after which the median fluorescence signal of the cells expressing empty pDRF1-GW plasmid was subtracted from all samples. Next, bleed-through was calculated using the ymTurquoise2 (ymTq2)-expressing strain, cells with an acceptor and donor fluorescence signal below 250 (arbitrary units) were discarded and FRET ratios were calculated for all remaining cells.

### gDNA extraction

Cells were scraped from a plate and resuspended in 100 μL Li-TE-SDS (100 mM lithium acetate, 10 mM Tris-HCl pH 7.5, 1 mM EDTA pH 8.0, 1% (v/v) SDS) and 1 μL of RNase A (10 mg/mL) was added to this mix. Cells were vortexed briefly and incubated at 65°C for 30 mins, with vortexing every 5 to 10 minutes. After incubation, the cells were centrifuged at 20000 g for three minutes. The supernatant was transferred to a new 1.5 mL microcentrifuge tube containing 300 μL ice-cold 100% ethanol. After gently mixing the tube, the DNA was centrifuged at 20000 g for 30 minutes at 4°C. The supernatant was removed and 100 μL of 70% (v/v) ethanol was added to the pallet. Next, the samples were centrifuged at 20000 g for 15 minutes at 4°C, supernatant was removed, and the residual ethanol was evaporated by air drying. The DNA pellet was resuspended in demiwater according to pellet size (20-50 μL).

### RNA extraction

W303-1A WT, Cyr1^K1876M^ and gpr1::LEU2 cells were grown as described with YNB medium containing 1% ethanol to mid-exponential phase. Next, cells were diluted in YNB medium containing 10 mM sucrose and grown overnight to mid exponential phase (OD_600_ 0.3-1) at 200 rpm and 30°C. Next, cells were centrifuged at 2000 g g for 3 minutes and frozen overnight at -20°C. Afterwards, cells were resuspended in 400 μL TES (10 mM Tris HCl, 1 mM EDTA and 0.5% (w/v) SDS at pH 7.8), 400 μL aquaphenol was added and incubated at 65°C for 3 minutes. Next, cells were put on ice for 5 minutes, spun down at 16000 g and 4°C, supernatant was transferred to a new tube and 400 μL aquaphenol was added and samples were spun down again at 2000 g for 10 minutes at 4°C after which the supernatant was transferred to a new tube and 400 μL chloroform was added and samples were vortexed. Next, cells were spun down at 16000 g and 4°C, supernatant was transferred and 160 μL of 2 M sodium acetate at pH5.2 was added. Next, DNA and RNA was precipitated by adding 5.2 mL of 100% ethanol and incubating the samples overnight at -20°C. The next day, the samples were centrifuged at 16000 g for 15 minutes at 4°C, supernatant was discarded and 1 mL of 70% ethanol was added. Finally, cells were centrifuged at 16000 g for 15 minutes at 4°C, supernatant was removed and the pellet was dissolved in 70 μL water.

Next, the samples were treated with DNase I (ThermoFisher Scientific) after which cDNA was synthesised using Maxima H Minus Reverse Transcriptase (ThermoFisher Scientific) according to the manufacturer’s protocol with RiboLock RNase Inhibitor (ThermoFisher Scientific) and Random Hexamers (ThermoFisher Scientific). DNA and RNA amounts were quantified using Qubit (ThermoFisher Scientific).

### qPCR

For the qPCR, primer efficiencies were determined by performing a qPCR with SYBR green (Applied Biosystems, Waltham, MA, United States) and a Applied Biosystems 7300 Real-Time PCR System using the manufacturer’s protocol with at least 4 concentrations of a 10-fold dilution series of gDNA and the primers listed in table 2 at 400 nM concentration, accompanied with the obtained primer efficiencies. Efficiencies were obtained in Excel (Microsoft, Redmond, WA, USA) by plotting Rt values versus ^10^Log of the DNA concentration after which a trendline was fitted and the obtained slope was used to determine the efficiencies according to equation 2.

**Table 2.**
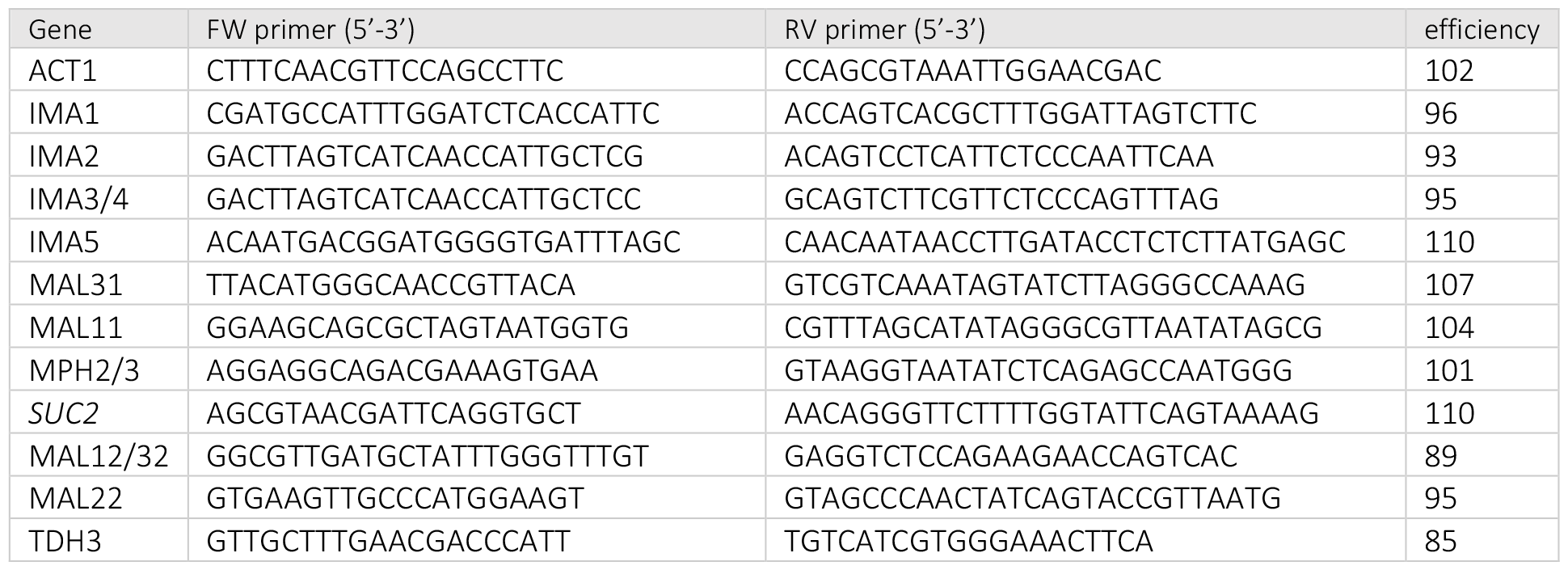
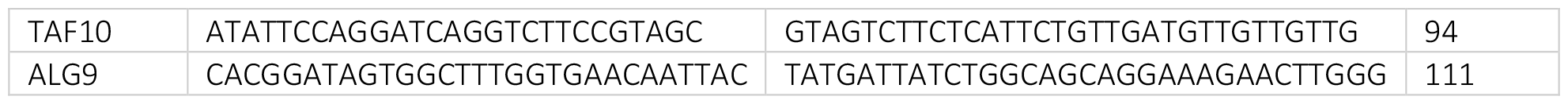
Used primers and the obtained efficiencies.

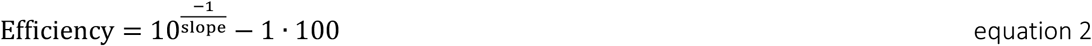

Next, qPCRs were performed using SYBR green using 5 ng/μL cDNA and 400 nM FW and RV primers according to manufacturer’s protocol. Next, Rt values were determined at a 0.07 threshold and expression ratios were determined using the Pfaffl Method^37^.

## Results

### cAMP signalling is more pronouncedly induced by sucrose than glucose

We hypothesise that Gpr1p functions as an extracellular sucrose sensor and therefore it should be properly activated by sucrose. By measuring population-based cAMP levels of cells growing on maltose, the affinity of the Gpr1p receptor for glucose was assessed and it was shown it reacts to glucose, but has a higher affinity for sucrose^35,38^. We re-assessed the affinity of the cAMP system to glucose and sucrose in ethanol-grown cells, using our recently published yEPAC sensor that measures cAMP levels in single yeast cells with high temporal resolution^38^. Fig. 2 shows the dose-response curves for glucose and sucrose, where the maximal peak height of the FRET response is plotted against the sugar concentration pulsed to ethanol-grown cells. Fitting the data with a Michaelis-Menten equation showed a significant better fit when fitting was grouped by the sugar pulsed (F-test, p<0.01). In contrast to previous results, we found a higher affinity for glucose and a higher maximal response for sucrose (Table 1). These results show that the cAMP signalling cascade can detect sucrose.

**Figure 2.**
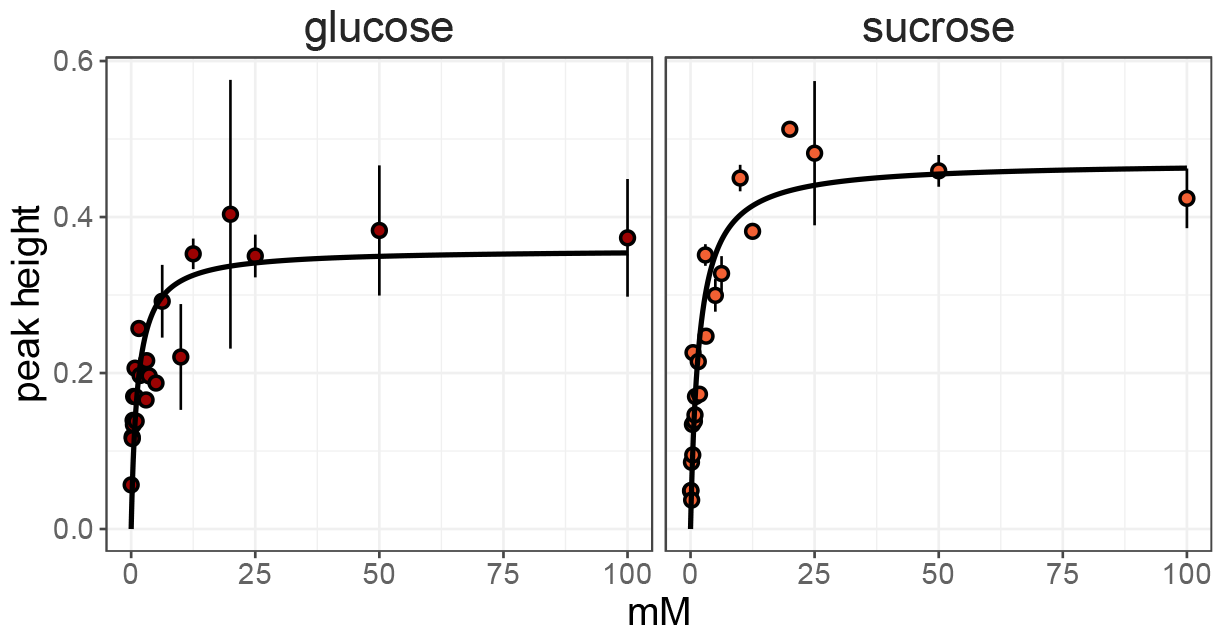
The cAMP system has a higher affinity for glucose, but a higher maximal response to sucrose. cAMP peak heights plotted against the amount of glucose or sucrose pulsed. Dots show mean cAMP peak height (baseline normalized), error bars indicate standard deviation, lines show Michaelis-Menten fit.

### cAMP signalling mutants show deteriorated responses to sucrose addition

Since we hypothesized that Gpr1p is involved in sucrose sensing, we tested whether the *Cyr1*^*K1876M*^ mutant (which has a mutation in the cAMP-producing enzyme adenylate cyclase) and the *gpr1::LEU2* knockout strain have impaired cAMP responses upon a sucrose transition. Ethanol pregrown cells were pulsed with either 100 mM glucose or sucrose and the cAMP levels were measured using the yEPAC FRET sensor (Figs. 3 and S1). Glucose addition still gave a cAMP peak in the mutant strains, but this response was largely absent during a transition to sucrose (maximal FRET increase of 20% and 21% for glucose and 8% and 10% for sucrose, for *Cyr1*^*K1876M*^ and *gpr1::LEU2* respectively). This suggests that cells lacking *GPR1* or having *Cyr1*^*K1876M*^ have more problems sensing sucrose than glucose, under these conditions. These results also indicate that the cAMP signalling cascade is important for sucrose signalling.

**Figure 3.**
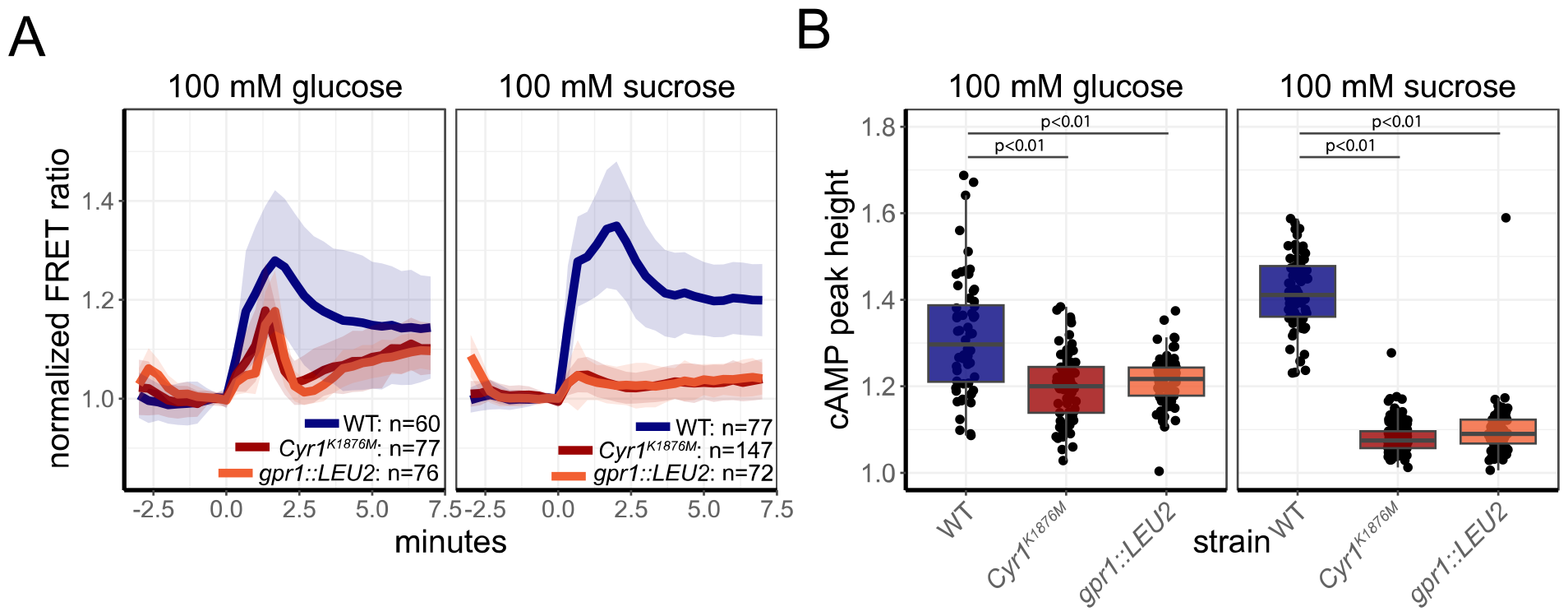
cAMP signalling mutants show a highly reduced cAMP response after sucrose addition. A) Normalized yEPAC FRET levels of W303-1A cells grown on 1% ethanol during a transition to either 100 mM glucose (left panel) or 100 mM sucrose (right panel). Lines show mean responses, line colour indicates the specific strain, shades indicate SD. B) Boxplot of the maximal FRET response after a 1% ethanol to 100 mM glucose (left panel) or 100 mM sucrose (right panel) transition for the 3 different W303-1A strains. Boxplot indicates median with quartiles; whiskers indicate largest and smallest observations at 1.5 times the interquartile range. Dots show the maximal response of each single cell. P-values of a Tukey-HSD test are shown between boxes.

### cAMP signalling mutants show a growth defect on sucrose, not on glucose

After having confirmed that the cAMP cascade indeed detects sucrose, we tested whether it also affects growth on sucrose. To do so, growth assays on different carbon sources were performed for the W303-1A WT and the *Cyr1*^*K1876M*^ strain (Fig. 4 and table 3). The strains were pre-grown on 1% (v/v) ethanol, washed twice and resuspended in YNB medium containing the necessary amino acids and 10 mM or 0.1% (v/v) of a carbon source. For growth on glucose and sucrose, the *gpr1::LEU2* strain was also included. Of the 12 carbon sources tested, only sucrose gave a significant reduction in growth rate between the strains (student t-test p-values shown in Fig. 4). The same holds for the lag phase (approximated as the time point after inocculation that gave the highest growth rate), which showed an increase in sucrose, although mannose also gave a significant difference. Mannose is known as an antagonist of the Gpr1p receptor^35^ and inhibition of this receptor in combination with a decrease cAMP production potentially increased lag time. The yield was significantly higher for both sucrose, fructose and glucose for only the *gpr1*Δ mutant. Although this work focussess mainly on the receptor-mediated cAMP signalling, we also assessed whether the other axis of cAMP signalling, the Ras2p axis, contributes specifically to sucrose signalling (Fig. S2). This was the case as the *ras2::KanMX* mutant showed a decreased growth rate and increased lag phase on sucrose but not on glucose. This indicates that impairing either one of the two signalling axes is sufficient to affect growth on sucrose.

**Table 3.** Fit parameters based on a Michaelis-Menten fit of the cAMP peak heights to glucose or sucrose. K_0.5_ is the sugar concentration giving half of the maximal response. Max response is the maximal increase (i.e., peak height) in baseline-normalized FRET of yEPAC after sugar addition. Fitting estimates ± standard error are given.

|  | Glucose | sucrose |
| --- | --- | --- |
| $K_{0.5}$ (mM) | 1.3 ($\pm 0.4$) | 1.7 ($\pm 0.3$) |
| Max | 0.36 ($\pm 0.02$) | 0.47 ($\pm 0.02$) |

**Figure 4.**
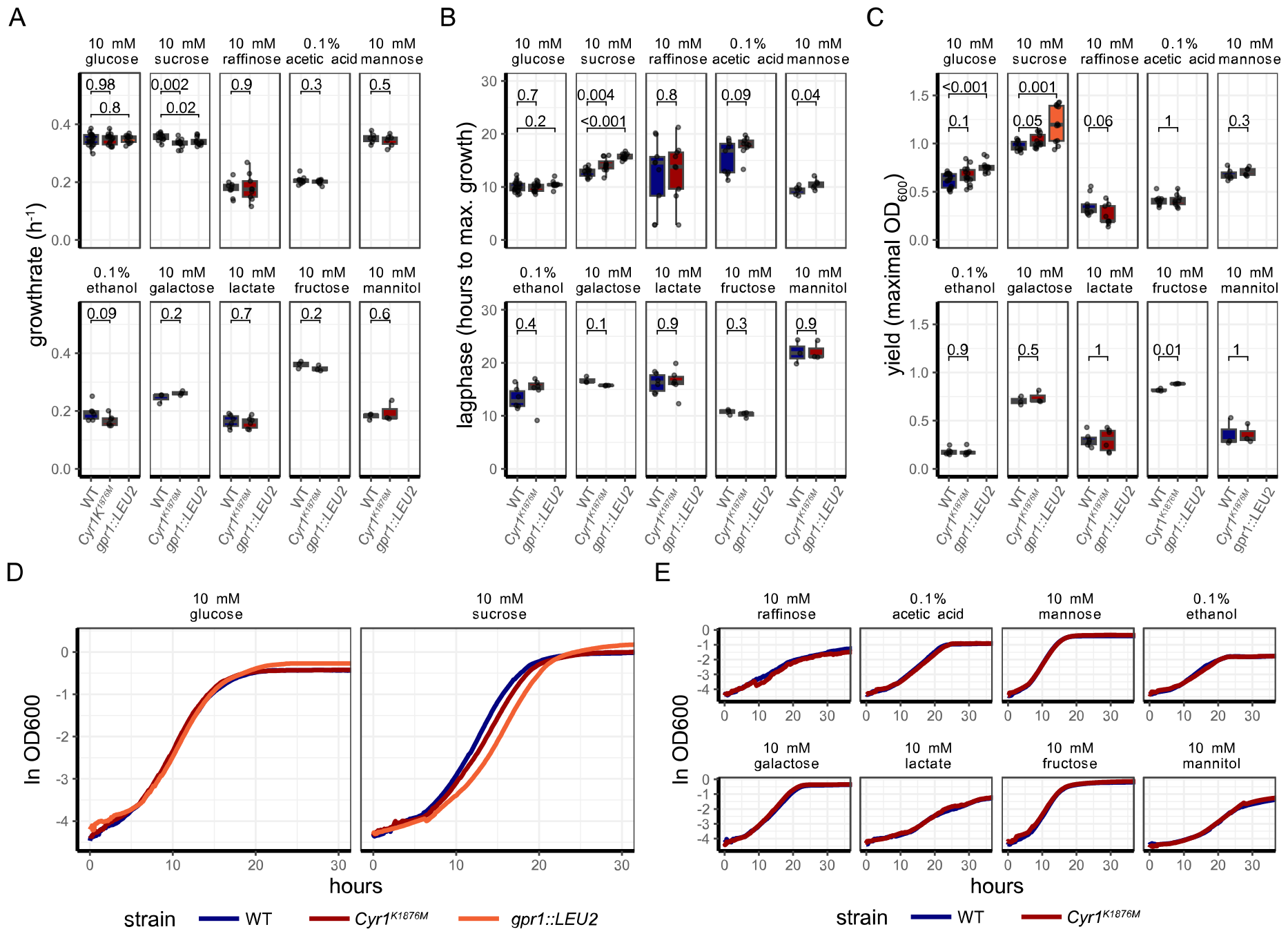
cAMP signalling mutants show an affected growth phenotype for sucrose only. A) Growth rates of W303-1A WT, Cyr1^K1876M^ and gpr1Δ mutants on various carbon sources. B) lag phase (i.e. time point at which cells obtain their maximal growth rate) of W303-1A WT, Cyr1^K1876M^ and gpr1Δ mutants on various carbon sources. C) Yield (i.e. maximal OD_600_ obtained during the growth assay) of W303-1A WT, Cyr1^K1876M^ and gpr1Δ mutants on various carbon sources. Boxplots depict median with quartiles; whiskers indicate largest and smallest observations at 1.5 times the interquartile range. Points show all replicates (technical and biological). D) Growth curves of W303-1A WT, Cyr1^K1876M^ and the gpr1Δ mutant on glucose and sucrose. E) Growth curves of W303-1A WT, Cyr1^K1876M^ and the gpr1Δ mutant on other carbon sources. Lines show the median ln OD_600_ for all replicates obtained, colour indicates strain. All data obtained from at least 2 biological replicates. Values depicted in panels A, B and C between boxes indicate p-values of student’s t-test.

Since cAMP signalling is also involved in stress signalling, we repeated the experiments but excluded the potential stress-inducing washing step during the inoculation of the growth assays. Furthermore, we included the *gpr1::URA3* strain to test whether the growth phenotype of the *gpr1::LEU2* is caused by potential prototrophial side effects. Again, we found decreased growth rates on sucrose for the *Cyr1*^*K1876M*^ and both *gpr1* knockouts, but not for glucose (Fig. 5 and table 4). Specifically sucrose gave an increased lag-phase for all strains but most significantly for the *gpr1* knockouts. Yield was increased for all mutant strains on both sucrose and glucose, but this effect was also more pronounced on sucrose (Fig. 5 and table 3).

**Table 4.**
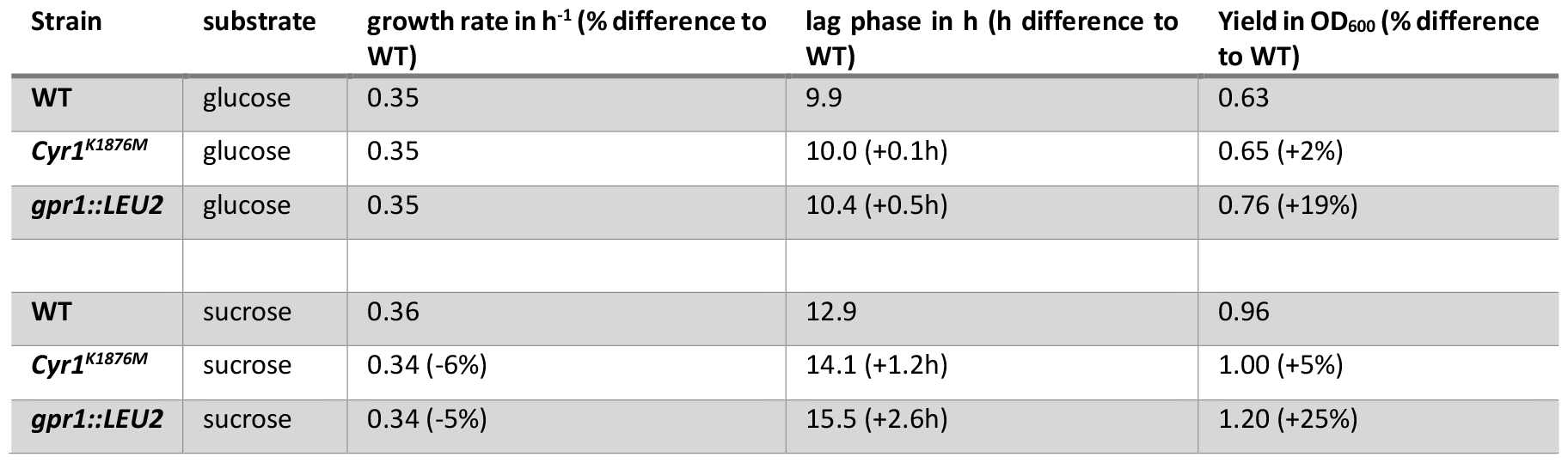
cAMP signalling mutants show impaired growth on sucrose. Growth characteristics of W303-1A WT, Cyr1^K1876M^ and gpr1::LEU2 pre-grown on 1% ethanol, washed in medium without carbon source, and directly transferred to 10 mM glucose and sucrose. Mean values and % difference to WT are shown, statistical significance between all strains and conditions can be found in figure 4.

**Figure 5.**
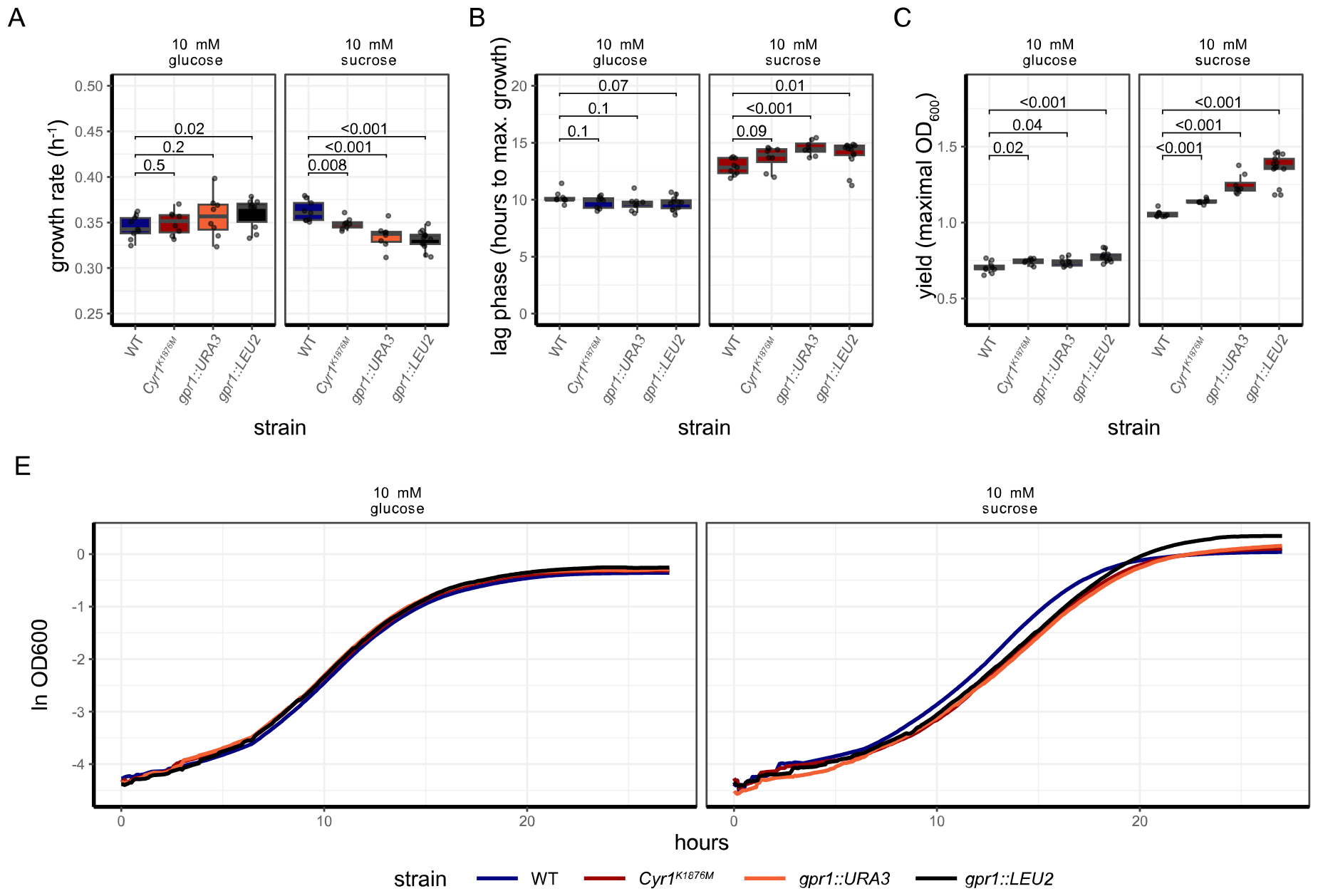
cAMP signalling mutants show affected growth also during a direct transition to sucrose without washing. A) Growth rates of W303-1A WT, Cyr1^K1876M^ and gpr1Δ mutants on various carbon sources. B) lag phase of WT and cAMP signalling mutants on glucose and sucrose C) Yield of the various strains on glucose and sucrose. D) Yield of the various strains on glucose and sucrose, corrected for the amount of carbon atoms each substrate contains. Boxplots depict median with quartiles; whiskers indicate largest and smallest observations at 1.5 times the interquartile range. Points show all replicates (technical and biological). E) Growth curves of W303-1A WT, Cyr1^K1876M^, gpr1::URA3 and gpr1::LEU2 on 10 mM glucose and sucrose. Lines show the median ln OD_600_ for all replicates obtained, colour indicates strain. Values depicted in panels A, B and C between boxes indicate p-values of student’s t-test. All data obtained from at least 2 biological replicates.

**Table 5.**
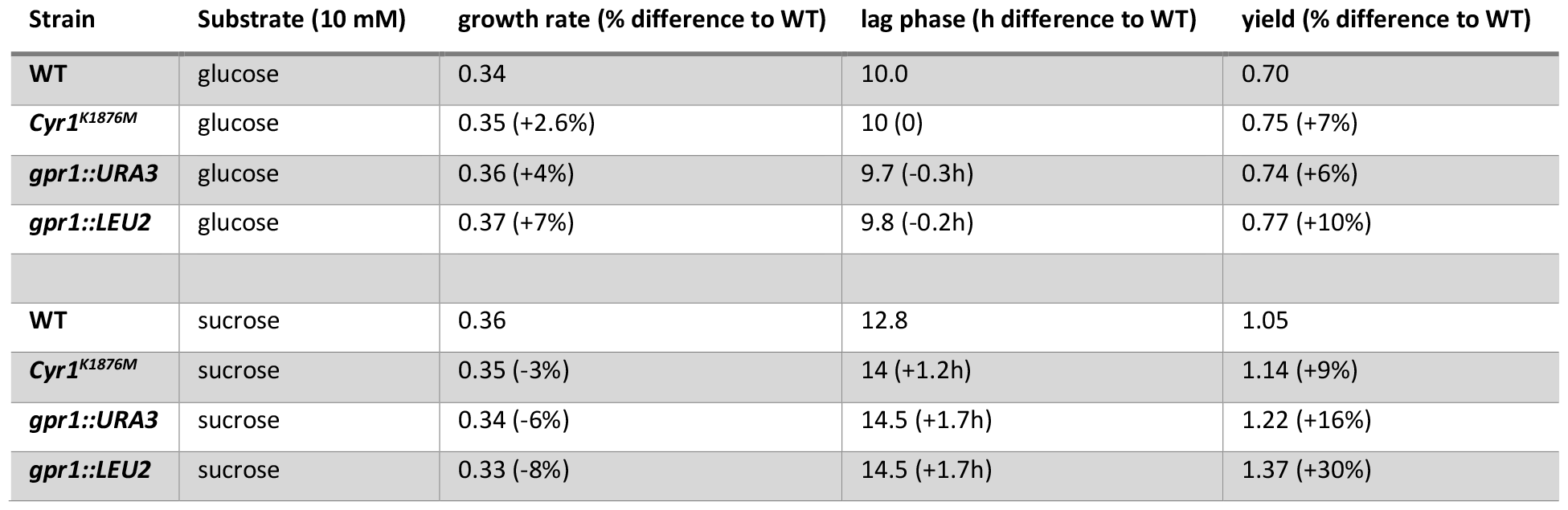
cAMP signalling mutants show more impaired growth on sucrose, also without washing. Growth characteristics of W303-1A WT, Cyr1^K1876M^, gpr1::URA3 and gpr1::LEU2 pre-grown on 1% ethanol and directly transferred to 10 mM glucose and sucrose. Mean values and % difference to WT are shown. Statistical significance between all strains and conditions can be found in figure 5.

In conclusion, in terms of growth phenotypes, the Gpr1-cAMP signalling mutants show impaired growth on specifically sucrose and no other carbon source.

### Impaired growth of the cAMP signalling mutants is not caused by decreased invertase activity

The metabolism of sucrose and glucose is identical, except for the hydrolysis of sucrose and contingent transport of this sugar. Thus, a decreased invertase activity is a potential cause of the mitigated growth phenotype of the cAMP signalling mutants on sucrose. To test this hypothesis, WT, *Cyr1*^*K1876M*^ and *gpr1::LEU2* cells were grown on 1% ethanol and inoculated into 100 mM sucrose medium and invertase activity was monitored using an invertase assay kit (Fig. 6A). In contrast to our hypothesis, an increased invertase activity was found for the *Cyr1*^*K1876M*^ mutant whereas the *gpr1::LEU2* mutant had similar invertase activity levels compared to WT. Thus, extracellular sucrose conversion is not at the basis of the altered growth characteristics of the cAMP-signalling mutants on sucrose. Potentially, the impaired growth found for the cAMP signalling mutants originates from other routes capable of consuming sucrose instead of invertase as a *suc2::URA3* mutant was still able to grow on sucrose (Fig. 6B-E). Other sucrose degrading pathways are the maltose transporters *MAL11/21/L31* and *MPH2/3* together with the isomaltoses *IMA1-5*^9,10^. Expression of these genes as well as *SUC2* was determined using qPCR. No significant difference in expression was found for these enzymes in all strains grown on 10 mM sucrose. Thus, the altered growth behaviour is not caused by altered invertase activity nor by altered expression levels of other sucrose metabolizing enzymes.

**Figure 6.**
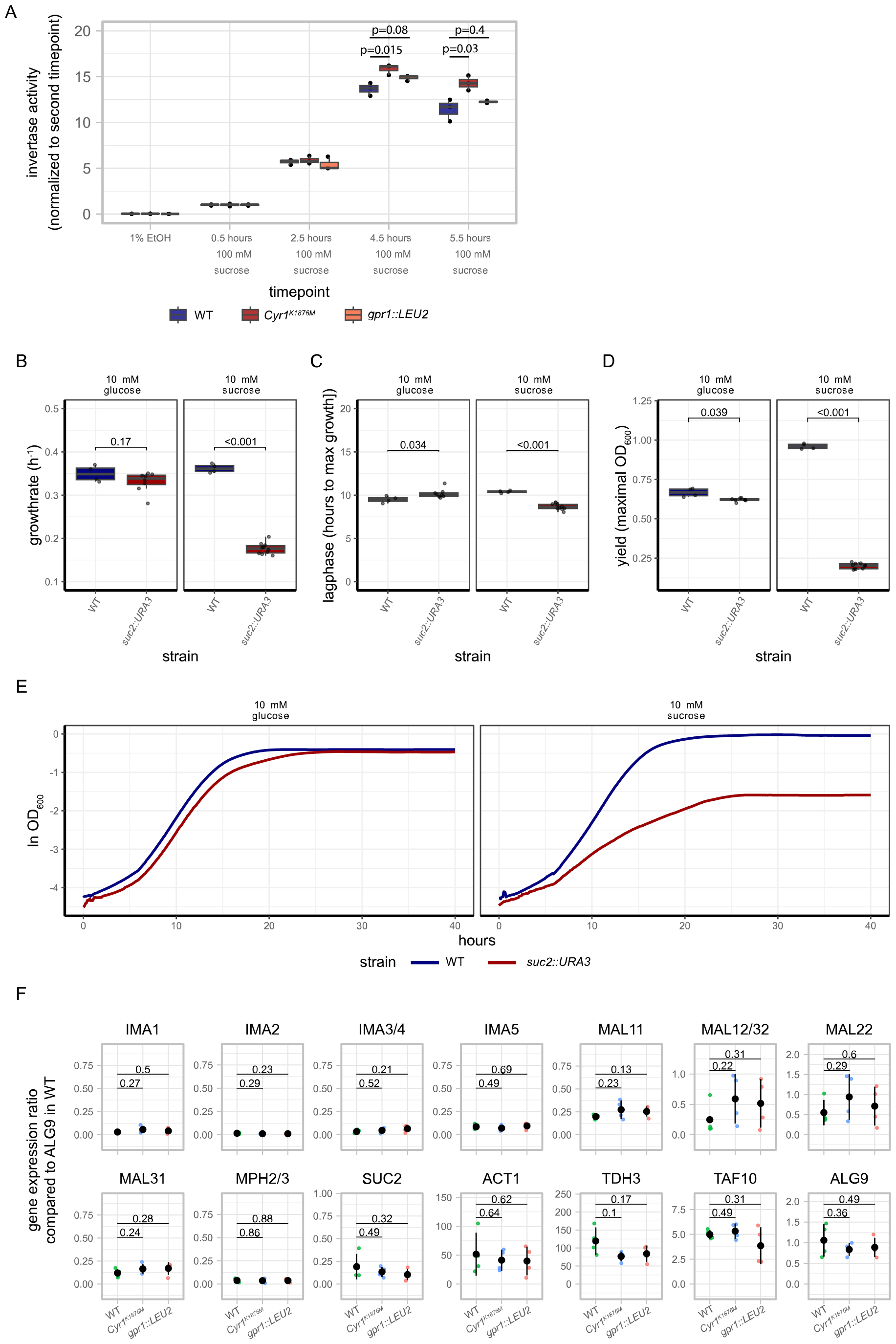
Invertase activity nor the maltose machinery lie at the basis of the affected growth phenotype. A) Invertase activity, normalized to the 0.5 hours 100 mM sucrose timepoint, is plotted against each timepoints. Each dot depicts a biological replicate, colour indicates strain. B) lag phase of WT and the SUC2 deletion mutant on sucrose C) Yield of the various strains on sucrose. E) Growth curves of W303-1A WT and the SUC2 deletion mutant on 10 mM glucose and sucrose. Lines show the median ln OD_600_ for all replicates obtained, colour indicates strain. Data obtained from at least 2 biological replicates. F) Gene expression levels of various sucrose metabolizing enzymes in W303-1A WT, Cyr1^K1876M^ and gpr1::LEU2 grown on 10 mM sucrose, normalized to ALG9 expression in W303-1A WT. Small dots represent biological replicates, colour indicates strain, black dots show mean, lines indicate SD. Data obtained from 4 biological replicates, p-values from student’s T-test are shown between the strains.

### Steady-state cAMP levels are not the cause of the altered growth phenotypes

The altered growth characteristics of the cAMP signalling mutants can either be caused by altered steady-state cAMP levels or the lack of the cAMP peak. Basal cAMP levels were measured using a FRET sensor-based reporter^38^ coupled to flow cytometry to assess whether this is altered between WT and the mutant strains (Fig. 7). FRET ratios of yEPAC were normalized to WT on 100 mM glucose and corrected for non-specific FRET changes by subtracting the FRET levels of the non-responsive yEPAC-R279L variant. FRET levels of yEPAC showed no significant differences compared to WT, which is in line with previous reports^34^. In conclusion, basal FRET levels are not the cause of the impaired growth behaviour of the cAMP signalling mutants. Altogether, we suggest that the lack of the short-term cAMP peak is possibly at the origin of the long-term altered growth phenotype on sucrose for these mutants.

**Figure 7.**
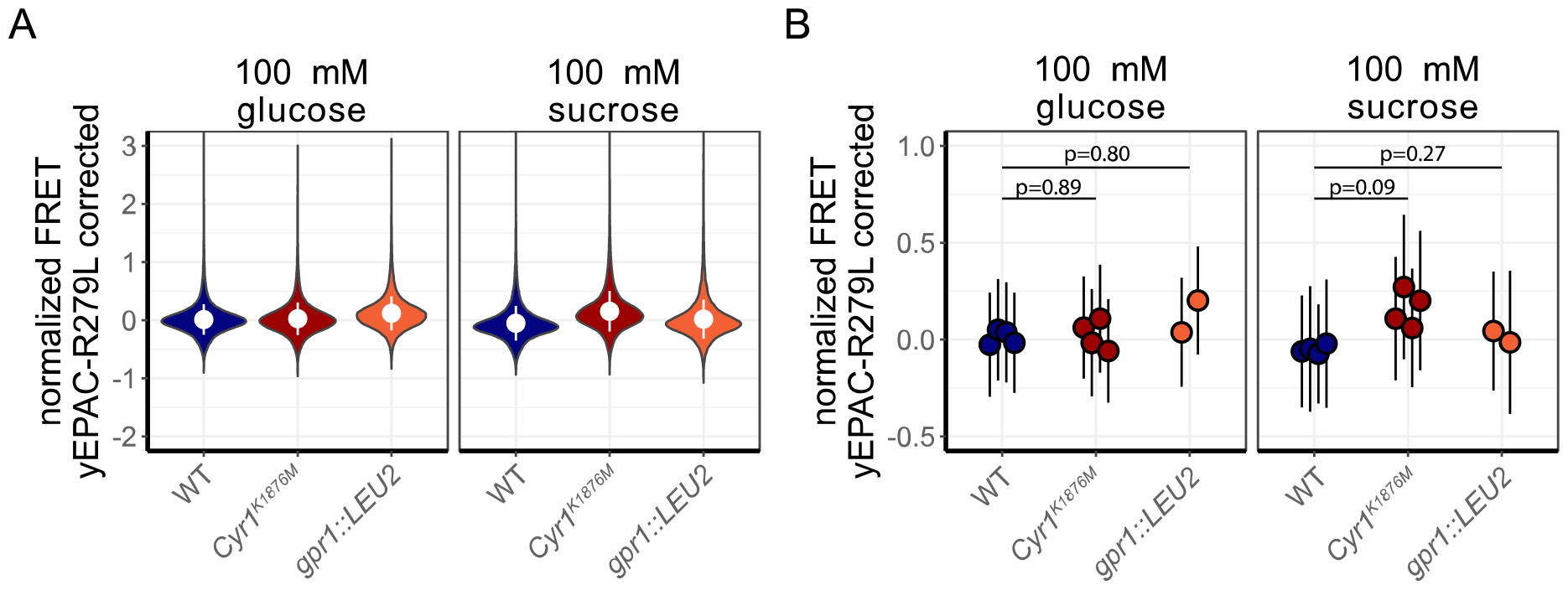
Basal cAMP levels measured using flow cytometry show no clear differences between cAMP signalling mutants and the WT strain. A) basal FRET levels yEPAC in W303-1A WT, Cyr1^K1876M^ and gpr1::LEU2 mutants grown on either 100 mM glucose or sucrose. Violin plot depict density of the FRET levels of all samples tested combined, dot depicts mean, vertical bars indicate SD. B) Basal FRET levels of the strains on either 100 mM glucose (left panel) or 100 mM sucrose (right panel) for each replicate. Each point depicts the mean of a biological replicate, error bars indicate SD, p-values of pairwise Wilcoxon rank sum tests are shown.

## Discussion

As sucrose is a prefered carbon source and a major industrial substrate for budding yeast, gaining insights in the regulation of sucrose metabolism and growth is of both fundamental and biotechnological importance. In this paper, we show several lines of evidence that suggest that cAMP signalling is important for transitions to growth on sucrose. The maximal cAMP response is higher for sucrose compared to glucose and the cAMP signalling mutants (*Cyr1*^*K1876M*^ and *gpr1Δ*) showed a highly abolished cAMP response when pulsed with sucrose, whereas glucose pulsing still gave a response. The glucose response in the signalling mutants strains is probably induced via the import and metabolism of glucose, which is less pronounced for a sudden sucrose transition as this needs to be converted extracellularly by invertase or metabolized by the maltose machinery (Fig. S3). The complete lack of a cAMP response for sucrose could mean that these cells miss the signal from Gpr1p about sucrose presence and therefore have more difficulties to adapt to this carbon resource.

This is indeed the case as the assessment of growth of the strains on various carbon sources show that only sucrose gives altered growth characteristics (with the increased yield on glucose for the signalling mutants being the only exception). These results were reconfirmed with growth assays without washing the cells. This implies that the cAMP signalling cascade is of importance for proper growth on sucrose for budding yeast. Of note, auxotrophic strains were used where certain markers were reintroduced to mutate components of the cAMP cascade. This can affect cellular physiology. Still, we are convincend that the found phenotypes are truly caused by the signalling cascades as we consistently see these phenotypes in various mutants, including the Cyr1^K1876M^ mutant which has no altered auxotrophies. The results are also in line with previous results showing that the Cen.PK strain, which has the Cyr1^K1876M^ mutation, has lower growth on sucrose^6,7^. The cAMP mutants strains in this study were still able to grow on sucrose at a high rate, indicating that cAMP signalling is not the sole signalling and regulating mechanism for sucrose metabolism or that the other modes of sucrose degradation degradation have a large, and possibly compensatory, influence on cellular growth. Indeed, *SUC2* and *IMA2* are known to be affected by other signalling pathways such as *Mig1*-*Snf1*^11,39–41^. Of note, we found that blocking either one of the two signalling axes is sufficient to impair growth on sucrose, which is in line with our previous findings that these two axes (i.e. startup of glycolysis and Gpr1p signalling) are both necessary to fully induce cAMP signalling in yeast^38^.

The impaired growth on sucrose is not caused by lowered extracellular invertase activity, a major mode of sucrose breakdown^13–17^. Likewise, gene expression levels of the maltose transporters and the isomaltoses are unaltered among the strains. In this study, the *Cyr1*^*K1876M*^ mutant even shows a slightly higher expression of invertase activity compared to the WT cells which is in line with a previous study^42^. The increased invertase activity could be attributed to glucose repression; cAMP signalling mutants showed a longer lag phase and a decreased growth rate, indicative of reduced glucose repression. *SUC2* is a commonly known glucose-repressed gene^43,44^. Perhaps the cAMP signalling mutants have decreased catabolite repression which increases invertase activity^43–45^. This would increase the higher flux of sucrose hydrolysis, but these strains are probably not optimally adapted to full fermentative growth and can therefore not benefit from the increased liberation of glucose and fructose from sucrose. This is an important trade-off for yeast cells; they can increase sucrose catabolism by decreasing catabolite repression to hydrolyse more sucrose per time unit, but this comes at the costs of correct metabolic rearrangement for a full fermentative (i.e. fully glucose-repressed) growth. cAMP potentially steers the cell optimally to solve this dilemma. The results reported here imply that the altered growth of the cAMP mutants strains is not caused by altered activity or expression of enzymes directly involved in sucrose metabolism, but rather by a more general metabolic adaptation issue.

A remaining question was whether the cAMP or steady-state cAMP levels affect the growth on sucrose. We found that basal cAMP levels are similar for the mutants and the WT strain. Thus, the cAMP peak is probably the main player to inducethe proper transition to sucrose metabolism, which means that this short-term response of a signalling cascade has an impact on subsequent cellular physiology and for multiple generations. When cAMP is not generated during the transition to sucrose, cells cannot fully and swiftly steer towards a fermentative, high-growth state and lose the competition with fellow organisms. This suggests that the cAMP peak is a short-term signal that tells the cell to make long-term investment in a fermentative, high-growth state. As cells residing in a high sucrose environment experience low levels of glucose and fructose (indicative for slow, respiratory growth), they need an independent signal reporting the presence of ample amounts of sugar resources to rapidly ferment on. We propose the cAMP signalling cascade provides this signal for sucrose.

In summary, we provide experimental evidence that support our hypothesis that the short-term dynamics of the cAMP signalling cascade, the cAMP peak, serve to sense sucrose and adapt yeast cells accordingly. In nature, where sucrose is a commonly used resource, the cAMP signalling cascade is likely an essential attribute for yeast cells to stay competitive. In industry, this signalling cascade can be exploited to either increase yield (i.e. by impairing cAMP signalling) or improve fermentation processes (i.e. by increasing the cAMP peak) where sucrose is used as a substrate.

## Data availability

All data and analysis scripts can be found online at DOI:10.17632/vzbsf357ds.1.

## Acknowledgements

We are grateful to Emile Zwering for helping with the Qubit system.

## Supplements

**Figure S1.**
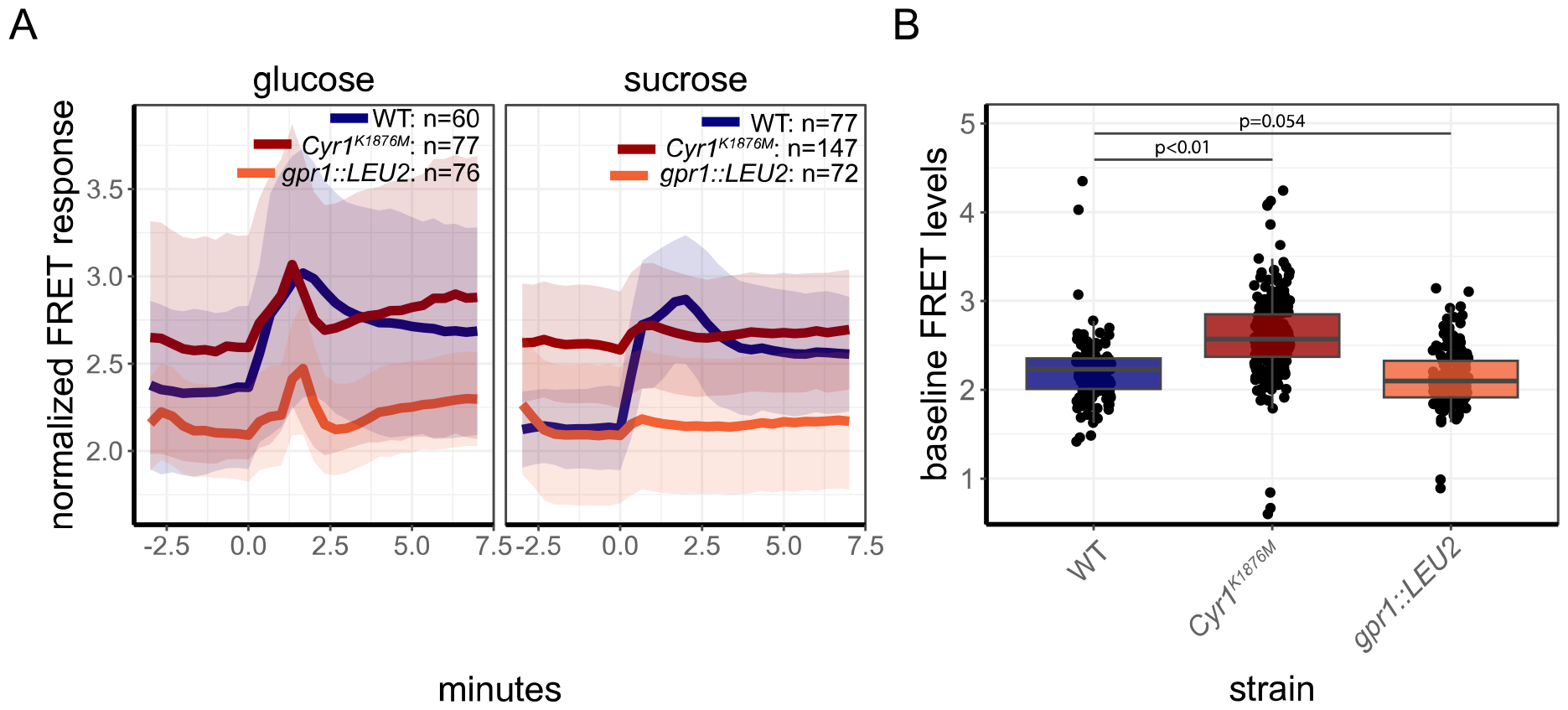
Raw yEPAC FRET responses after a sugar addition. A) yEPAC FRET levels of W303-1A cells grown on 1% ethanol during a transition to either 100 mM glucose (left panel) or 100 mM sucrose (right panel). Lines show mean responses, line colour indicates the specific strain, shades indicate SD. B) Boxplot of the baseline FRET levels for the various strains. Boxplot indicates median with quartiles; whiskers indicate largest and smallest observations at 1.5 times the interquartile range. Dots show the maximal response of each single cell. P-values of a Tukey-HSD test are shown.

**Figure S2.**
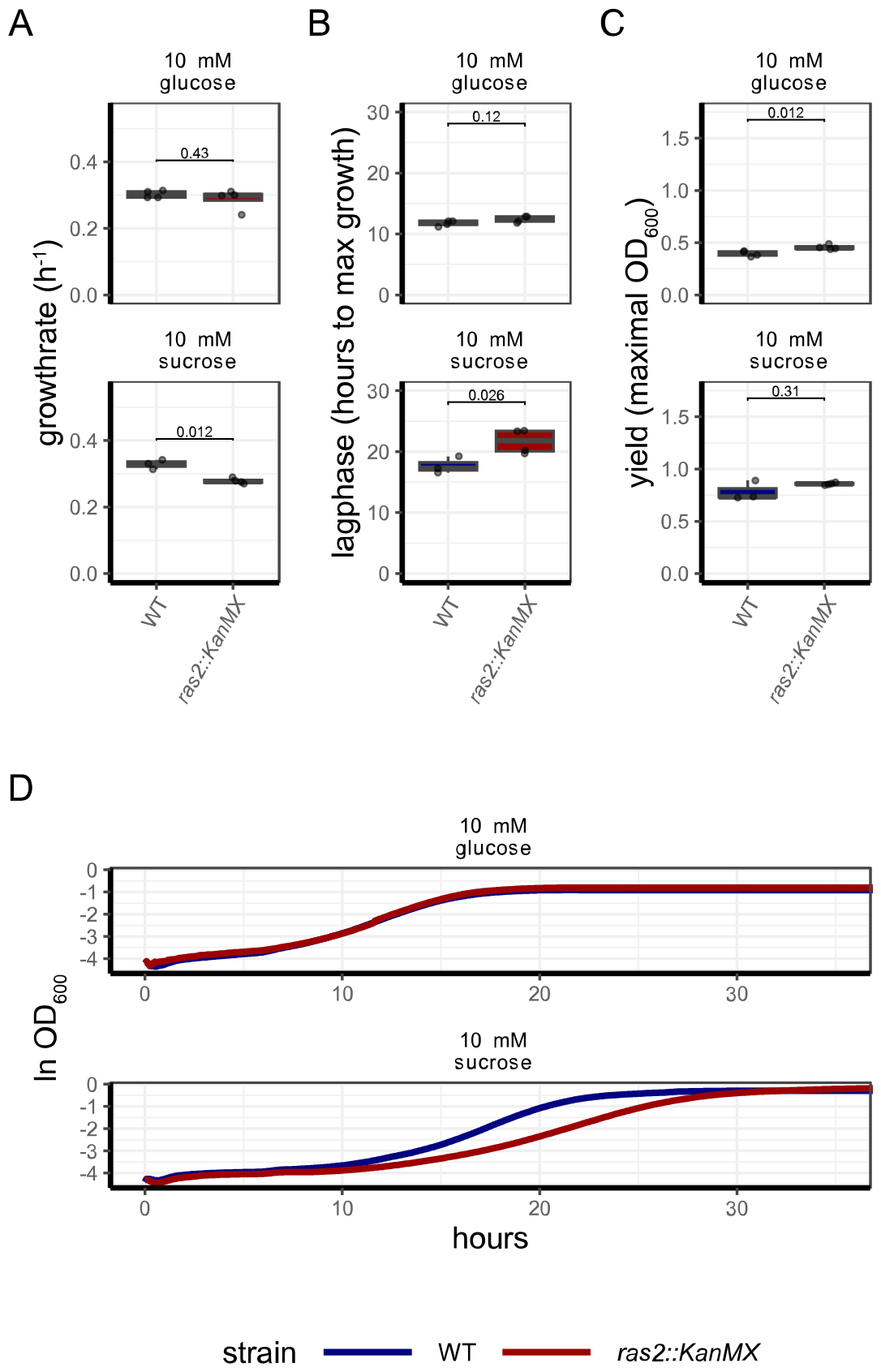
The ras2 deletion mutant shows a growth effect on sucrose. A) Growth rates of W303-1A WT and ras2::KanMX on either 10 mM glucose or sucrose. B) lag phase (i.e. time point at which cells obtain their maximal growth rate) of W303-1A WT and ras2::KanMX on either 10 mM glucose or sucrose. C) Yield (i.e. maximal OD_600_ obtained during the growth assay) of W303-1A WT and ras2::KanMX on either 10 mM glucose or sucrose. Boxplots depict median with quartiles; whiskers indicate largest and smallest observations at 1.5 times the interquartile range. Points show all replicates (technical and biological). D) Growth curves of W303-1A WT and ras2::KanMX on either 10 mM glucose or sucrose on various carbon sources. Lines show the median ln OD_600_ for all replicates obtained, colour indicates strain. All data obtained from at least 2 biological replicates.

**Table S1.** The ras2 deletion mutant shows a growth defect on sucrose. Growth characteristics of W303-1A WT and ras2::KanMX pre-grown on 1% ethanol, washed in medium without carbon source, and directly transferred to 10 mM glucose and sucrose. Mean values and % difference to WT are shown.

| Strain | substrate | growth rate (% difference to WT) | lag phase (h difference to WT) | yield (% difference to WT) |
| --- | --- | --- | --- | --- |
| WT | glucose | 0.30 | 11.9 | 0.40 |
| *ras2::KanMX* | glucose | 0.30 | 12.5 (+0.6h) | 0.45 (+12%) |
| WT | sucrose | 0.33 | 17.3 | 0.73 |
| *ras2::KanMX* | sucrose | 0.28 (-16%) | 21.8 (+4.5h) | 0.85 (+16%) |

**Figure S3.**
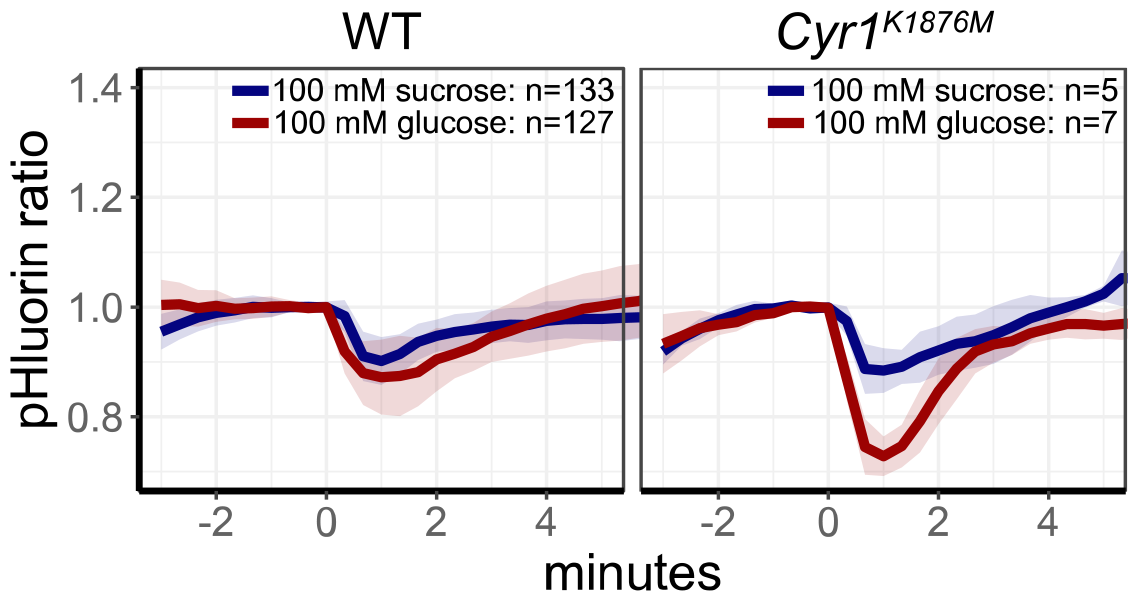
pH dynamics, measured by pHluorin, as a measure for glycolytic startup. The pH response of WT or Cyr1^K1876M^ mutated W303-1A cells to either glucose or sucrose. Lines show the mean response, line colours indicate the pulsed sugar, shades indicate SD.

